# In Vitro Efficacy of “Essential Iodine Drops” Against Severe Acute Respiratory Syndrome-Coronavirus 2 (SARS-CoV-2)

**DOI:** 10.1101/2020.11.07.370726

**Authors:** Köntös Zoltán

## Abstract

**Background:** Aerosolization of respiratory droplets is considered the main route of coronavirus disease 2019 (COVID-19). Therefore, reducing the viral load of Severe Acute Respiratory Syndrome-Coronavirus 2 (SARS-CoV-2) shed via respiratory droplets is potentially an ideal strategy to prevent the spread of the pandemic. The *in vitro* virucidal activity of intranasal Povidone-Iodine (PVP-I) has been demonstrated recently to reduce SARS-CoV-2 viral titres. This study evaluated the virucidal activity of the aqueous solution of Iodine-V (a clathrate complex formed by elemental iodine and fulvic acid) as in Essential Iodine Drops (EID) with 200 μg elemental iodine/ml content against SARS-CoV-2 to ascertain whether it is a better alternative to PVP-I.

**Methods:** SARS-CoV-2 (USAWA1/2020 strain) virus stock was prepared by infecting Vero 76 cells (ATCC CRL-1587) until cytopathic effect (CPE). The virucidal activity of EID against SARS-CoV-2 was tested in three dilutions (1:1; 2:1 and 3:1) in triplicates by incubating at room temperature (22 ± 2°C) for either 60 or 90 seconds. The surviving viruses from each sample were quantified by a standard end-point dilution assay.

**Results:** EID (200 μg iodine/ml) after exposure for 60 and 90 seconds was compared to controls. In both cases, the viral titre was reduced by 99% (LRV 2.0). The 1:1 dilution of EID with virus reduced SARS-CoV-2 virus from 31,623 cell culture infectious dose 50% (CCCID_50_) to 316 CCID_50_ within 90 seconds.

**Conclusion:** Substantial reductions in LRV by Iodine-V in EID confirmed the activity of EID against SARS-CoV-2 in vitro, demonstrating that Iodine-V in EID is effective at inactivating the virus *in vitro* and therefore suggesting its potential application intranasally to reduce SARS-CoV-2 transmission from known or suspected COVID-19 patients.

## Introduction

The global pandemic of Coronavirus disease 2019 (COVID-19) caused by the Severe Acute Respiratory Syndrome-Coronavirus 2 (SARS-CoV-2) was first identified in the city of Wuhan, China during late 2019 (1). It has since spread rapidly globally, with over 26 million confirmed cases to date and over 865000 deaths. The rapid spread of this novel SARS-CoV-2 virus is enhanced by the shading of viral particles from infected or symptomatic individuals through bodily secretions, especially respiratory droplets including saliva, and nasal fluid. It has been demonstrated that tiny droplets of 0-10 μm diameter produced during speech-making, sneezing, coughing, and panting contain SARS-CoV-2 viral particles (2), which remained substantially viable and infectious in aerosols for up to 3 hours, similar to SARS-CoV-1 (3). Thus, it is presumed that the most effective infection mechanism is the aerosolization of viral particles in respiratory droplets (4), which is easily inoculated directly into the airway during breathing or indirectly by contact transfer via contaminated hands (5). The aerosolized microscopic infectious viral particles can linger in the air circulating in poorly ventilated rooms or spaces long enough to cause multiple infections to individuals who inhale them (6). A raft of recommended preventive measures including wearing face masks, eye protection, washing hands and keeping social distance on > 1 m have been demonstrated to be effective in reducing COVID-19 transmission (7). The goal of face masks is to reduce the transmission of respiratory droplets from infected individuals and shield non-infected individuals from transmitted droplets. Face masks as vital Personal Protective Equipment (PPE) against COVID-19 disease is not fully accepted and has remained controversial from perspectives of attitude, effectiveness, and necessity (8).

In the initial stage of COVID-19 disease, SARS-CoV-2 viral titres of > 107/ml in saliva and nasal mucous have been reported; and therefore, reduction of these titres should help to reduce disease transmission from known COVID-19 patients to individual who is uncomfortable using masks (9). This can be achieved using oral and nasal sprays with antiviral activity against SARS-CoV-2 virus. Intranasal saline sprays containing elemental iodine as an active ingredient has been used for nasal moisturizing and prevention and/or treatment of sinusitis or rhinitis (10). Iodine-containing intranasal/oral moisturizing saline sprays are being explored as drug agents against coronaviruses. Their potential use has been supported by a recent clinical safety trial that demonstrated that intranasal Povidone-Iodine (PVP-I; 1.25%) spray had no adverse effects for up to five months and therefore, could be used against coronaviruses (SARS-CoV-1/2 and Middle East Respiratory Syndrome - MERS) (11). Regarding the antiviral efficacy of iodine-containing nasal/oral sprays, nasal and oral formulations containing PVP-I have been reported to inactive SARS-Cov-2 *in vitro* (12, 13). Thus, PVP-I oral antiseptic is potentially efficacious in reducing the transmission risk of coronaviruses in dental practice (13). SARS-CoV-2 is internalized into the host cells via two viral receptors of host cell infection: Angiotensin-Converting Enzyme 2 (ACE2) and CD147 (a highly-glycosylated transmembrane protein) by binding to them using virus spike proteins (SP) (14). The mechanism of action of PVP-I mouth rinse/nasal spray involves targeting of the host’s ACE2 for inhibition. It inactivates the activity of Haemagglutinin Esterase (HE), a fifth important structural protein of beta-coronaviruses and diminishes the ACE2 receptors in lymphocytes by promoting their absorption from host epithelial tissue (14). This reduces the concentration of SARS-CoV-1/2 shed in saliva and nasal fluid.

Enthused by promising findings from a recent study by Pelletier and colleagues (12), the present study was interested in Essential Iodine Drops (EID) for oral/nasal decontaminant in known or suspected cases of COVID-19 as a potentially better alternative to PVP-I. Iodine-V is a novel complex derivative of fulvic acid in a clathrate complex by elemental iodine (I_2_) molecule. Essential Iodine Drops (EID) is an aqueous solution of Iodine-V containing 200 μg elemental iodine/ml. EID is currently used as an oral dietary supplement to support a healthy thyroid function and is non-toxic, non-allergic and lends itself to prolonged continuous use if kept within the daily recommended upper limit of 1100 μg. Its potential for use in protecting against the transmission of SARS-CoV-2 virus during elective surgical procedures in ophthalmology, otolaryngology and dental has been demonstrated recently using PVP-I solutions (12). Whilst steps are already being made to make Essential Iodine Drops (EID) available for nasal administration, the present study was conducted to ascertain whether the EID solution is effective in deactivating SARS-CoV-2 *in vitro*, as demonstrated with PVP-I. Therefore, this research aims to evaluate the *in vitro* virucidal efficacy of Essential Iodine Drops (EID) on SARS-CoV-2 to ascertain whether it is a better alternative to PVP-I.

## Materials and Methods

The present study tested the virucidal efficacy of Essential Iodine Drops (EID) against SARS-CoV-2 using similar materials and protocol by Pelletier and colleagues (12). Importantly, the present study was conducted in Biosafety Level 3 (BSL-3) laboratories at The Institute for Antiviral Research at Utah State University, USU (Logan, UT) following established Standard Operating Procedures approved by the USU Biohazards Committee.

### Cell lines and virus

Before testing, a virus stock of SARS-CoV-2 (USAWA1/2020 strain) sourced from the CDC following its original isolation from an infected patient in Washington state, USA (WA-1 strain BEI #NR-52281 strain), was prepared according to Pelletier and colleagues (12). The virus stock was prepared by infecting Vero 76 cells (ATCC CRL-1587) until cytopathic effect (CPE) was visible at two days postinoculation. The Vero 76 cells were cultured in Minimal Essential Medium (MEM) (Quality Biological), supplemented with 2% (v/v) Fetal Bovine Serum (Sigma), and 50 μg/ml gentamicin (Gemini Bio-products)(12).

### Drug testing

The test drug was Essential Iodine Drops (EID), obtained from IOI Investment Zrt. (Budapest, Hungary). It is the aqueous solution of Iodine-V. The Iodine-V is a water-soluble elemental iodine in fulvic acid clathrate complex. Physical and chemical properties of Iodine-V are as follows: iodine content as determined by Inductively Coupled Plasma Mass Spectrometry (ICP-MS) was 10.01%; the melting point was 157°C, the maximum ultra-violate/visible (UV/VIS) absorption in the water at 340 nm was 96 μg/ml); the spectral signature of Iodine-V as analysed by Fourier-Transform Infrared Spectra (FTIR) in KBr pastille contained the following wavenumbers (v; cm-1) of key signals: 3404, 2927, 2359, 2341, 1718, 1635, 1558, 1418, 1383, 1151, 1076, 1024, 668. The melting point was determined in an SRS 100 OptiMelt apparatus (Stanford Research Systems, Sunnyvale, CA, US). Ultraviolet was measured in a DLab SP1100 system. FT-IR was measured in FTIR650 (Labfreez Instruments Group Co., Ltd.).

The virucidal activity of Essential Iodine Drops against SARS-CoV-2 was tested in three dilutions; 1:1; 2:1 and 3:1. The original concentration of Essential Iodine Drops as supplied by IOI Investment Zrt was 200 μg elemental iodine/ml. This was subsequently diluted with SARS-CoV-2 virus solution to 1:1 for 90 seconds, 2:1 for 60 seconds and 3:1 for 60 seconds.

### Virucidal Assay

Briefly, the three dilutions of Essential Iodine Drops (EID) containing SARS-CoV-2 virus solution (1:1; 2:1 and 3:1) were tested in triplicates for virucidal activity as described by Pelletier and colleagues (12). The undiluted drug as supplied (without virus solution) in two tubes was used as toxicity and neutralization controls. Ethanol (70%) was used as the positive control while water was used as a virus control. The test solution and virus were incubated together at room temperature (22 ± 2°C) for either 60 seconds and 90 seconds and the solution were then neutralized by a 1/10 dilution in MEM with FBS (2%) and gentamicin (50μg/ml).

### Virus Quantification

The survived viruses from each sample were quantified by a standard end-point dilution assay according to Pelletier and colleagues (12). Briefly, neutralized samples were pooled and serially diluted using eight log dilutions in the test medium. Subsequently, 100 μL of each dilution was plated into quadruplicate wells of 96-well, plates containing 80-90% confluent Vero 76 cells. The toxicity controls were added to an additional 4 wells of Vero 76 cells and 2 of those wells at each dilution were infected with the virus to serve as neutralization controls, ensuring that the residual sample in the titre assay plate did not inhibit growth and detection of the surviving virus. Plates were incubated (37 ± 2°C with 35% CO2) for 5 days. Each well was then scored for the presence or absence of the infectious virus. The titres were measured using a standard endpoint dilution 50% cell culture infectious dose (CCID50) assay calculated using the Reed-Muench (1938) equation and the log reduction value (LRV) of each compound compared to the negative (water) control was calculated(12).

## Results

Infectious SARS-CoV-2 viruses were quantified by endpoint dilution virus titration on Vero cells and the results were expressed as log_10_ cell culture of 50% infectious dose (log_10_ CCID_50_). Virus titres and log reduction value (LRV) of SARS-CoV-2 when incubated with a single concentration of the test drug for each time point are shown in Table 1. After 90 seconds of incubation, 1:1 EID reduced viral titre by 99% from 4.5 log_10_ CCID_50_/0.1 ml to 2.5 log_10_ CCID_50_/0.1 ml giving an LRV of 2.0. It reduced the virus from 31,623 CCID_50_ to 316 CCID_50_ per 0.1ml. After 60 seconds, 3:1 EID reduced viral titre by 99% from 4.0 log_10_ CCID_50_/0.1 ml to 2.0 log_10_ CCID_50_/0.1 ml giving an LRV of 2.0. No cytotoxicity was observed in any of the test wells and both positive and neutralization controls performed as expected.

**Table 1:** Virucidal efficacy of Essential Iodine Drops (EID) against SARS-CoV-2 60 and 90 seconds incubation with the virus at 22 ± 2°C.

| Test Article | EID concentrations | Incubation Time | Virus Titer<sup>a</sup> | LRV<sup>b</sup> |
| --- | --- | --- | --- | --- |
| EID | 66 % | 60s | 2.3 | 1.7 |
| EID | 75 % | 60s | 2.0 | 2.0 |
| Ethanol (70%) | 50 % | 60s | <0.67 | >3.33 |
| Virus Control | n.a. | 60s | 4.0 | n.a. |
| EID | 50 % | 90s | 2.5 | 2.0 |
| Ethanol (70%) | 50 % | 90s | <0.67 | >4.33 |
| Virus Control | n.a. | 90s | 4.5 | n.a |

<sup>a</sup> Log<sub>10</sub> CCID<sub>50</sub> of virus per 0.1 ml. The assay lower limit of detection is 0.67 Log<sub>10</sub> CCID<sub>50</sub>/0.1 ml.
<sup>b</sup> LRV (log reduction value) is the reduction of virus compared to the virus control. n.a.=not applicable.

## Discussion

Iodine is established as having a broad-spectrum antimicrobial activity against bacterial, viral, fungal and protozoal pathogens and has been used as an antiseptic for the prevention of infection and the treatment of wounds for decades. PVP-I currently has the best antiviral activity against SARS-CoV and MERS-CoV (9), and its virucidal *in vitro* activity against the novel, SARS-CoV-2 virus has been demonstrated recently (12). This paper aimed to demonstrate that the novel Iodine-V in EID as an active virucidal ingredient is as efficacious as PVP-I and potentially safer. In vitro virucidal assay in the present study has indeed demonstrated that 75% and 50% of Essential Iodine Drops (EID) solution reduced SARS-CoV-2 virus titre after 60 seconds and 90 seconds of incubation by an LRV of 2.0 (99%). These findings indicate that Iodine-V in EID lends itself as a drug with dual benefits, a mineral supplement to maintain a healthy thyroid functioning and reduce the transmission of SARS-CoV-2 virus. When applied orally or intranasally, Iodine-V in EID is potentially beneficial against the transmission of SARS-CoV-2 from COVID-19 patients. This means that the health benefits of EID are superior to that of PVP-I or Lugol’s iodine (iodine and potassium iodide in water alone forming mostly triiodide).

Although PVP-I was reported to be non-cytotoxic (12), this formulation may cause serious rashes that are similar to chemical burns observed in a few rare cases. It has been implicated in late-onset allergic contact dermatitis when used as a pre-operative antiseptic in oral and maxillofacial surgery (15). This has been attributed to the presence of free iodine (I_2_), which has a strong oxidizing effect on the skin or mucosa, hence, the reported allergic reaction. Free iodine concentrations in PVP-I (1.25% dilution) and Lugol’s iodine are regarded as low and safe. The virucidal activity of EID was lower than an LRV of 4.6 for PVP-I (99.99%) as demonstrated by Pelletier et al. (12), however, 1-5% PVP-I containing 1000-5000 μg/ml iodine compared to 200 μg/ml iodine in EID. Elemental iodine in Iodine-V within the fulvic acid clathrate complex forming a solid stable material that can be easily formulated into a tablet form for slow, extended release of elemental iodine. This means that even EID tablet formulation would be less likely to cause iodine-induced allergies in oral and nasal mucosae. Furthermore, EID is formulated with Iodine-V without excipients unlike PVP-I and therefore, has a potentially better virucidal activity against SARS-CoV-2 virus. Furthermore, PVP-I excipient has been reported, in rare cases, to induced immediate (type 1) hypersensitivity reactions in children (16). Therefore, EID can be considered a safer alternative to PVP-I as it can be used in children.

Unlike PVP-I, which is routinely used in surgical settings as a pre-operative antiseptic in oral and maxillofacial surgery (15), EID is currently used in the general populace as a mineral supplement and can be readily accepted as a drug to prevent the spread of the COVID-19 pandemic. Being already on the market as a mineral supplement to help people with iodine deficiency, data on its potential adverse effects are already known or available and therefore, safe when used routinely orally or intranasally in known or suspected COVID-19 patients.

## Conclusion

Iodine-V (currently only available in EID formulation) inactivated 99% of SARS-CoV-2 after 60 and 90 seconds; quite similar to Povidone-Iodine (PVP-I) (99.99% at 60 seconds) reported elsewhere. Being excipient-free, this data also suggests that Iodine-V in EID is likely to have better stability and an enhanced potency *in vivo* when compared with PVP-I. Therefore, Iodine-V offers an advantage as a nasal or oral antiseptic to reduce viral transmission from known or suspected COVID-19 patients. With the increasing home-based care approach for COVID-19 patients, the risk of transmitting SARS-CoV-2 virus among family members and the immediate community can be reduced using intranasal/oral Iodine-V, which has the potential to remove or inactivate shed virus from respiratory secretions of known or suspected COVID-19 cases.

## Abbreviations

SARS-CoV-2: Severe Acute Respiratory Syndrome-Coronavirus 2
COVID-19: Coronavirus Disease 2019
PVP-I: Povidone-Iodine
EID: Essential Iodine Drops
Iodine-V: iodine - fulvic acid clathrate complex
Vero 76: ATCC CRL-1587
CPE: Cytopathic Effect
CCID_50_: cell culture infectious dose 50% assay

## Declarations

### -Ethics approval and consent to participate

This *in vitro* study used SARS-CoV-2 (USAWA1/2020 strain) sourced from the CDC, which obtained it anonymously from an infected patient in Washington State, USA. Therefore, ethics approval and patient consent are not applicable in this section.

### -Consent to publication

Not applicable.

### - Availability of data and material

The data supporting study findings can be found in a separate supplemental material that can be accessed by reaching out to the corresponding author, Dr. Köntös Zoltán, through his

### -Authors’ contributions

Dr. Köntös Zoltán is the sole researcher who coordinated laboratory analysis and wrote the manuscript.

## Acknowledgements

I am particularly grateful for the service given by Jonna B. Westover with The Institute for Antiviral Research at Utah State University.

